# CREST: A Cortical Resting-State EEG Spatial Transformer for Chronic Pain Inference

**DOI:** 10.64898/2026.08.25.747119

**Authors:** Yasha Iravantchi, Edward Lannon, Sean Mackey

## Abstract

Chronic pain mechanisms are complex, spanning multiple brain regions and networks. We ask whether resting brain activity carries a readout of that state. From a few minutes of resting-state electroencephalography (EEG), we generate a spectrogram to represent how each region of the cortex oscillates across frequency and time and pass it through `CREST` (<u>C</u>ortical <u>R</u>esting-state <u>E</u>EG <u>S</u>patial <u>T</u>ransformer): a frozen image-recognition network that reads each region as an image—here, a spectrogram—paired with a graph model that weighs the 56 cortical regions together to classify chronic-pain status. Across 125 people (74 with chronic pain, 51 healthy controls), evaluated through a leave-one-subject-out cross-validation, `CREST` separates the two groups with an area under the receiver operating characteristic curve (AUROC) = 0.782 (permutation *p <* 0.005). Control experiments implicate each person’s individual alpha rhythm.

**Clinical relevance:** A resting-state EEG readout of chronic MSK pain could clarify pathophysiology and inform treatment.

## I. Introduction

Chronic musculoskeletal (MSK) pain is among the most prevalent and disabling conditions in medicine, yet its underlying mechanisms remain poorly understood. The neuroplasticity driven by chronic MSK pain extends beyond nociceptive pathways to distributed circuits spanning the major brain networks—the default mode, salience, and central executive networks. Most human recordings, however, sample a single network at a time, leaving inter-network interactions unmeasured and brain models of pain correspondingly incomplete. Whether distributed resting activity across these networks contains information sufficient to distinguish chronic MSK pain from healthy status remains an open question, one whose answer could inform both basic understanding and treatment development.

Resting-state electroencephalography (EEG) is an appealing sensing modality. It is inexpensive and noninvasive, and it captures the ongoing electrical rhythms of the cortex, several of which (e.g., alpha and beta) are altered in chronic pain [1], [2]. Resting-state eyes-closed (EC) and eyes-open (EO) engage distinct patterns of brain communication, biased toward interoceptive and exteroceptive processing, respectively. Furthermore, prior EEG studies of pain mostly rely on single-network, hand-picked spectral or connectivity features and small samples, and are often not evaluated in a way that guarantees the model has learned pain rather than memorized individuals. These hand-crafted approaches may be challenging to reimplement or may not generalize; we take a deliberately generic approach built from core signal-processing and machine-learning components.

We convert resting source-localized EEG into per-region spectrograms, images of how each cortical region’s rhythms vary across frequency and time, and classify chronic-pain status with `CREST`, a *Cortical* (activity mapped onto the cortex itself, i.e., source space) *Resting-state EEG Spatial Transformer* that reasons over the anatomical layout of the cortex.

This paper makes the following contributions:

1. `CREST`, a method that takes any per-region signal rendered as an image, encodes each with a frozen ImageNet-pretrained network, and reasons over the cortex with a hemisphere-aware graph transformer;
2. instantiated on alpha-anchored spectrograms, this purely spectral representation decodes chronic-pain status at an AUROC of 0.78 under leave-one-subject-out cross-validation (*N* = 125);
3. ablations locate that performance: removing the pretrained backbone, the cortical graph, or the positional encoding each degrades the result, with pretraining the dominant factor;
4. anchoring to each person’s own alpha peak removes the age-confounded absolute position of alpha, and control experiments implicate that individualized band.

## II. Related Work

### Mechanistic Understanding of Chronic Pain

Despite decades of effort, the search for the mechanistic underpinnings of MSK pain has spanned imaging, autonomic signals, and electrophysiology without converging on a routine measure. Functional MRI (fMRI) signatures separate acute pain states but are costly and impractical for routine or repeated use; autonomic measures (e.g., heart-rate variability, skin conductance) are cheap but nonspecific. Resting-state EEG has drawn sustained interest because chronic pain is accompanied by changes in cortical rhythms and connectivity [1]–[3]. Furthermore, reported classifiers tend to use small cohorts and hand-crafted features, and few establish that performance holds for previously unseen individuals.

### Machine Learning for Clinical Biomarkers

Clinical biomarker discovery uses machine learning on high-dimensional physiological data, where the principal constraints are small sample sizes and the risk of learning confounds instead of the target; subject-independent evaluation and confound analysis are the accepted safeguards.

### Transformers and Spatial Models for Brain Data

Transformers [4] and graph neural networks have been applied to brain data by treating anatomical regions as nodes and their spatial or functional relationships as edges. In parallel, time-frequency representations of physiological signals have been decoded with convolutional networks pretrained on natural images [5], [6]. `CREST` combines these threads: a pretrained image backbone as a per-region encoder, and a graph transformer over a cortical parcellation as the spatial reasoner.

## III. Materials and Methods

We describe `CREST` through a four-stage pipeline: 1) cohort acquisition; 2) spectrogram construction; 3) per-region encoding; and 4) spatial aggregation. We now describe each stage in further detail.

### Cohort Acquisition

We study 125 adults with complete resting-state recordings: 74 with chronic pain (i.e., persisting beyond 3 months; group SC, Subject Chronic) and 51 healthy controls (group SH, Subject Healthy), a 59/41 split. The groups differ in age (SC 57.1 *±* 15.0 years, SH 40.6 *±* 14.3 years) and sex (SC 79.7% female, SH 43.1% female), which we address in the cohort confound analysis below. Data were collected with informed consent under an approved IRB protocol. Each participant contributed a resting-state EEG recording in two conditions, eyes-closed (EC) and eyes-open (EO). Continuous 250 Hz EEG was source-localized to a 56-region, hemisphere-labeled cortical parcellation.

### EEG Spectrograms

For each of the 56 cortical regions we compute a spectrogram from its source-localized time course, using a 10 s window advanced in 1 s steps over a 120 s segment, across 2–45 Hz, then interpolating the time axis to 224 columns. Each region contributes an eyes-closed and an eyes-open spectrogram. The two are encoded separately and combined in feature space (`CREST` fuses this way whenever a region carries more than one condition) by concatenation, absolute difference, and elementwise product.

Before the filterbank we remove the aperiodic (1*/f*) background, itself age-dependent [7], by subtracting a per-frame log-log linear fit and keeping the positive residual; filterbank outputs are log-compressed to form the image. EfficientNet-B0, our per-region encoder, has a native resolution of 224*×* 224. The alpha rhythm is the roughly 8–12 Hz oscillation that dominates the resting EEG and is strongest with the eyes closed. Its exact peak frequency differs from person to person, near 10 Hz on average but drifting 2 *±* Hz; this per-person peak is the individual alpha frequency (IAF), a stable trait consistent within a person across sessions [8] that also declines with age [9]. We estimate each participant’s IAF from their eyes-closed spectrum [10] and align the filterbank to it: a triangular filterbank packs its centres inside IAF 4 *±* Hz and thins out above it, so the alpha band fills a broad stripe of the image (≈13 rows/Hz inside the window vs. ≈ 2 above; Fig. 2); for the 4 of 125 participants lacking a clear alpha peak we fall back to 10 Hz. We anchor on alpha because: 1) cortical rhythms in and around the alpha band are altered in chronic pain [1], [2]; 2) the IAF has been proposed as a marker of pain sensitivity [3]; and 3) the thalamocortical-dysrhythmia hypothesis, one influential but debated account, links chronic pain to a shift of resting rhythms toward lower frequencies [11]. Anchoring to each person’s own peak removes the absolute position of alpha, the part most confounded with age, and emphasizes the shape of activity around it instead. The incremental-over-demographics test guards against what remains.

**Fig. 1.**
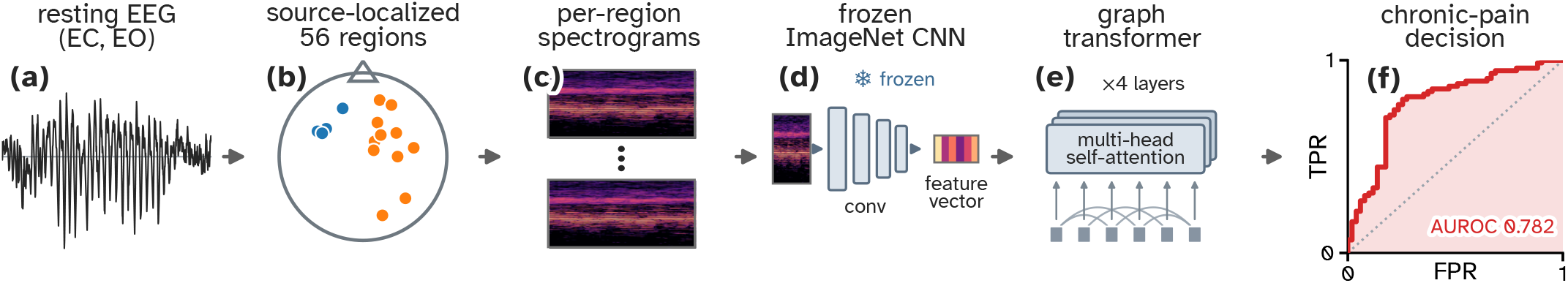
The CREST pipeline. Resting-state EEG is (**a**) recorded (eyes closed and open) and (**b**) source-localized to 56 cortical regions; (**c**) each region’s rhythms become an alpha-anchored spectrogram; (**d**) an image network pretrained on photographs, never retrained on EEG, summarizes each one; (**e**) the 56 summaries are compared across the cortex; and (**f**) the model reports a chronic-pain probability (ROC curve, AUROC 0.782, leave-one-subject-out, *N* = 125).

**Fig. 2.**
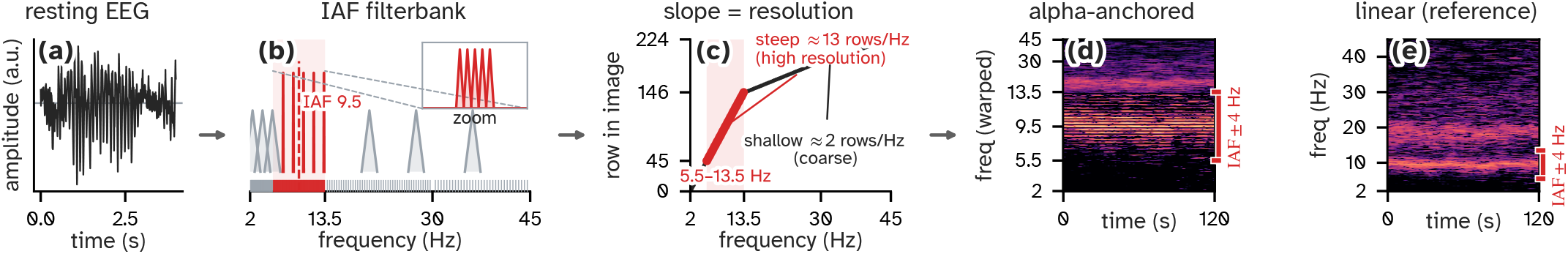
Spending the image’s resolution on each person’s own alpha rhythm. **(a)** Source-localized EEG from one cortical region. **(b)** A filterbank anchored to this participant’s alpha peak (IAF, 9.5 Hz) puts 101 of its 224 filters inside IAF *±* 4 Hz, concentrating the image’s detail there. **(c)** The resulting map from frequency to image row is steep below and inside that window and flat above it (≈ 2 rows/Hz). **(d)** The image `CREST` reads: alpha fills a broad band of the 224 *×* 224 canvas. **(e)** The same signal with uniform frequency spacing, for reference: alpha is squeezed into a thin stripe. The red bracket marks IAF *±* 4 Hz in both. Re-anchoring this window away from each participant’s own peak degrades decoding (see Table II).

### Spectrogram Encoding via EfficientNet

A spectrogram’s discriminative content tends to lie in local time-frequency texture (the density and regularity of banding) rather than in object-like shapes, so we reasoned that a network trained on natural images, which are known to favor local texture over global shape [12], might transfer well as a general-purpose spectrogram encoder without any EEG-specific training. We therefore embed each region’s spectrogram with a frozen, ImageNet-pretrained EfficientNet-B0 [5], [6], global-average-pooling its output to a per-region feature vector (Fig. 3). Freezing the backbone keeps the method simple; the ablations below show that a randomly initialized backbone collapses below chance.

**TABLE 1.** Cohort demographics.

| Group | $n$ | Age (yr) | Female (%) |
| --- | --- | --- | --- |
| Chronic pain (SC) | 74 | $57.1 \pm 15.0$ | 79.7 |
| Healthy (SH) | 51 | $40.6 \pm 14.3$ | 43.1 |

**TABLE II.**
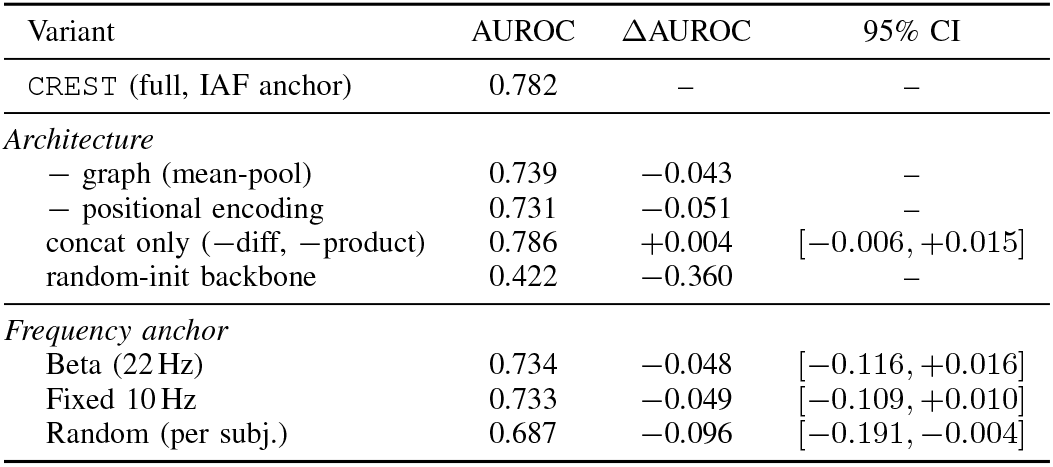
Ablations, baselines, and frequency-anchor controls (LOSO, 5-seed; paired ΔAUROC VS. full CREST).

| Variant | AUROC | ΔAUROC | 95% CI |
| --- | --- | --- | --- |
| CREST (full, IAF anchor) | 0.782 | — | — |
| *Architecture* |  |  |  |
| — graph (mean-pool) | 0.739 | −0.043 | — |
| — positional encoding | 0.731 | −0.051 | — |
| concat only (−diff, −product) | 0.786 | +0.004 | [−0.006, +0.015] |
| random-init backbone | 0.422 | −0.360 | — |
| *Frequency anchor* |  |  |  |
| Beta (22 Hz) | 0.734 | −0.048 | [−0.116, +0.016] |
| Fixed 10 Hz | 0.733 | −0.049 | [−0.109, +0.010] |
| Random (per subj.) | 0.687 | −0.096 | [−0.191, −0.004] |

**Fig. 3.**
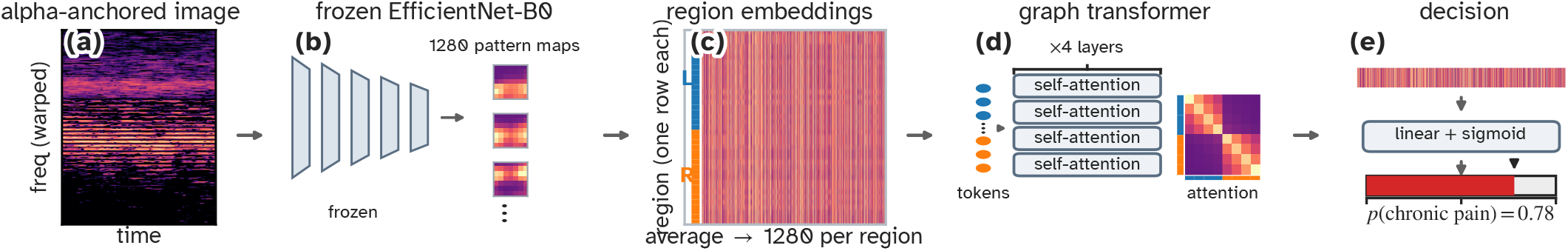
From one region’s spectrogram to a decision. **(a)** One region’s alpha-anchored spectrogram is treated as an image. **(b)** An off-the-shelf image-recognition network (EfficientNet-B0), trained on photographs and never retrained on EEG, reports how strongly each of 1280 learned visual patterns appears in it; 3 are shown. **(c)** Averaging each pattern’s response across the image summarizes the region as 1280 numbers. The identical network is applied to every region, giving one row per region, tagged by hemisphere. **(d)** The regions are then compared with one another, each pair weighted by cortical distance (a four-layer graph transformer; pairings shown are illustrative). **(e)** The regions are combined into a single summary and converted to a chronic-pain probability. An untrained network of identical size collapses below chance (see Table II).

### The CREST Architecture

`CREST` takes any per-region image as input; the alpha-anchored spectrogram above is the instantiation we evaluate here. The 56 per-region feature vectors form the nodes of a four-layer graph transformer [4] with 4 attention heads and model dimension 384 (the 1280-d per-region embeddings are projected to 384 before attention) (Fig. 3). Region identity enters through a fixed sinusoidal positional encoding over the parcellation’s 7 × 8 row/column layout, mirrored so bilateral homologs share a position. Attention is further biased by a learned 56 *×* 56 matrix initialized to negative cortical distance, so nearby and homologous regions start more strongly coupled; this bias is what makes the transformer a graph over the cortex. A learned class token attends over the regions alongside them, and its output representation is passed to a linear-plus-sigmoid head that emits chronic-pain probability. Figure 1 shows the full pipeline.

### Evaluation Approaches

We evaluate with strict leave-one-subject-out (LOSO) cross-validation: every fold holds out one participant entirely, so no person appears in both training and testing. Results are averaged over 5 random seeds as an ensemble. Statistical significance uses a subject-shuffle permutation null. We report the AUROC with subject-level percentile-bootstrap confidence intervals and the area under the precision-recall curve against the cohort base rate.

## IV. Results and Discussion

### Classification Performance

Under strict LOSO cross-validation (*N* = 125; 74 chronic-pain, 51 healthy), `CREST` separates the two groups at an AUROC of 0.782 (95% bootstrap CI 0.69–0.87 over held-out subjects) and an area under the precision-recall curve of 0.820 against a base rate of 0.592. We report the AUROC of the seed-averaged predictions; the 5 seeds vary only initialization and training order, never the folds. The observed AUROC exceeds all *B* = 200 subject-shuffle permutation nulls (null mean 0.42, below 0.50 due to the 59/41 class imbalance; maximum 0.63), each of which re-runs the entire pipeline on relabeled subjects (*p* = (1+0)*/*(1+200) *<* 0.005), so it is not an artifact of the finite sample. Overall, the separation is well above chance and stable across seeds.

### Ablation Evaluations

The headline number alone does not say which component earns it, so we ablate each under an identical evaluation (Table II). The dominant factor by far is ImageNet pretraining: replacing the frozen backbone with a randomly initialized one of identical architecture collapses AUROC from 0.782 to 0.422, below chance, so domain-general pretrained features—not the network’s capacity—carry essentially the entire signal. On top of that representation the spatial model adds a smaller but consistent margin: a mean-pooled baseline that discards the region graph entirely still reaches 0.739, and the full graph transformer recovers 0.782 (a paired +0.043). That margin rests on the positional encoding: removing it drops the graph to 0.731, no better than pooling. Conversely, concatenation alone is statistically indistinguishable from the full fusion (0.786; paired ΔAUROC +0.004, 95% CI [*−* 0.006, +*−*0.015]). A gradient-boosted-tree (XGBoost) baseline on the same features reaches 0.673. Thus, pretraining supplies the representation, and the cortical graph supplies a smaller spatial increment on top of it.

### Signal Importance and Interpretation

Moving our frequency emphasis away from each person’s alpha peak should degrade the decoder if it reads genuine alpha physiology, and it does. Anchoring the emphasis to the individual alpha peak yields AUROC 0.782; re-anchoring the same resolution budget onto the beta band drops it to 0.734, fixing it at a common 10 Hz to 0.733, and scattering it to a random frequency per subject to 0.687 (Table II). At this sample size only the drop against a random per-subject anchor is statistically significant (paired ΔAUROC 0.096, 95% CI [*−*0.191, *−* 0.004]); the beta and fixed-10 Hz drops are of similar magnitude but their paired confidence intervals cross zero. A band-nonspecific component remains, since the random anchor still exceeds chance, but anchoring to each person’s own peak adds a measurable increment over a randomly placed band. Ultimately, part of the discriminative signal appears tied to each participant’s own alpha rhythm.

### Cohort Confound Evaluation

The chronic-pain and healthy groups differ in age and sex (Table I), so we ask whether `CREST` merely reflects demographics. A linear fit of the `CREST` score on age and sex explains only 10.7% of its variance (*R*2 = 0.107). Adding the score to a logistic model of age and sex improves the fit significantly over demographics alone (likelihood-ratio 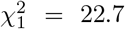, *p* = 1.9 *×* 10^*−*6^), so `CREST` captures pain-related brain activity over and above a demographic proxy.

### Discussion

Multi-network brain modeling can help elucidate the brain changes associated with chronic pain symptoms. A reproducible EEG correlate of chronic pain could complement self-report by providing an additional neurophysiological phenotype, particularly for longitudinal studies and treatment-response research. `CREST` does not establish an objective ground truth for pain and should not be interpreted as validating or invalidating an individual’s report; evaluation in independent clinical cohorts and longitudinal within-person studies is necessary before clinical application.

Methodologically, `CREST` is a compact recipe for small-sample clinical EEG. Because the backbone is frozen and pretrained, the approach needs little labeled data and no EEG-specific representation learning. Inference is correspondingly cheap—a single forward pass through a frozen EfficientNet-B0 per region plus a four-layer transformer—though the source-localization step upstream remains the principal requirement on a deployment site.

The study has clear limits. It is single-site with a modest, demographically imbalanced sample, and it decodes pain status at a single time point rather than tracking change. CNS-active medications common in chronic pain also alter resting spectra and are not in the confound model. Source coverage also varies by recording: regions without a reconstructed dipole are omitted, so the graph operates on the populated subset of the parcellation. The alpha localization is reported as a property of the signal; we make no mechanistic claim about what the network computes. External validation and longitudinal testing are the natural next steps.

## V. Conclusion

`CREST` decodes chronic-pain status from resting-state EEG spectrograms at an AUROC of 0.78 under strict LOSO evaluation, with each pipeline component shown to contribute and the discriminative signal implicating each person’s individual alpha rhythm. Because the backbone is frozen and pretrained, the approach generalizes to any per-region signal rendered as an image. It offers a compact, reusable method and a candidate EEG correlate of a condition still measured almost entirely by self-report.

